# Physiological and transcriptomic responses to N-deficiency and ammonium: nitrate shift in *Fugacium kawagutii* (Symbiodiniaceae)

**DOI:** 10.1101/2020.05.04.077719

**Authors:** Tangcheng Li, Xibei Chen, Senjie Lin

## Abstract

Symbiodiniaceae are the source of essential coral symbionts of reef building corals. The growth and density of endosymbiotic Symbodiniaceae within the coral host is highly dependent on nutrient availability, yet little is known about how Symbiodiniaceae respond to the dynamics of the nutrients, including switch between different chemical forms and changes in abundance. In this study, we investigated physiological, cytometric, and transcriptomic responses in *Fugacium kawagutii* to nitrogen (N)-nutrient deficiency and different chemical N forms (nitrate and ammonium) in batch culture conditions. We mainly found that ammonium was consumed faster than nitrate when provided separately, and was preferentially utilized over nitrate when both nitrogen compounds were supplied at 1:2, 1:1 and 2:1 molarity ratios. Besides, N-deficiency caused decreases in growth, energy production, antioxidative capacity and investment in photosynthate transport but increased energy consumption. Growing on ammonium produced a similar cell yield as nitrate, but with a decreased investment in nutrient transport and assimilation. These all have important implications of N nutrient to support symbiosis in coral ecosystem, especially regarding ammonium. In addition, by integrating our current results with previous data, we identified ten highly and stably expressed genes as candidate reference genes, which will be potentially useful for gene expression studies in the future.

## 1. Introduction

The symbiotic relationship between scleractinian corals and their endosymbiotic photosynthetic algae from the family of Symbiodiniaceae forms the foundation of coral holobionts and shallow-water coral reefs (Radecker et al., 2015). In this association, the phototrophic Symbiodiniaceae provide photosynthates that may meet up to 95% of the coral’ energy requirement; in exchange, the corals provide shelter and inorganic nutrients to their algal partners (Muscatine et al., 1997; Yellowlees, et al., 2008). However, this relationship is delicate and susceptible when the environment condition alters. The loss of the dinoflagellate symbionts and/or their pigment from the coral in response to environment stress can lead to coral bleaching and ultimately death of the coral and destruction of the reef (Hoegh-Guldberg, 1999). On the contrary, excessively high density of the symbionts can also harm the symbiotic relationship and even turn the relationship into parasitism under certain environment conditions (Thrall et al., 2007). For this mutualistic association to prosper, corals need to regulate their symbiont density both on a seasonal basis and in response to local environmental conditions (Fitt et al., 2000; Cunning et al., 2015). And one way to regulate this is through controlling nitrogen (N)-nutrient availability (Radecker et al., 2015).

The highly efficient uptake and recycling of nutrients by coral reef organisms endow tropical reef-building corals with the exceptional ability to flourish in nutrient-poor environments (Muscatine et al, 1977). In particular, N-cycling in the coral holobionts may be crucial for the coral-algae symbiosis. The availability of N nutrients shows strong seasonal and diel variations and can be affected by anthropogenic activities (D’Angelo and Wiedenmann, 2013). Anthropogenic eutrophication often results in not only an increase of N and phosphorus (P) availability but it also usually modifies the N: P stoichiometry, all of which are critical for maintaining the algal community structure (Brodie et al., 2011; Lapointe et al., 2019). Several studies reported increased algal density in response to elevated concentrations of dissolved inorganic nitrogen (DIN) in the water (Muscatine et al., 1989; Fabricius et al., 2005), indicating a strong influence of the external N levels on the proliferating rates of Symbiodiniaceae and other algae in the environment, the latter of which may cause deleterious effects on coral health.

In the natural ecosystems, various forms of DIN exist, mainly including nitrate and ammonium. When nitrate is taken up, it needs to be reduced first to nitrite and then to ammonium before it can be assimilated into amino acids, and nitrate reduction requires energy, so ammonium is assumed the preferred form of nitrogen nutrient for Symbiodiniaceae (Shepherd et al., 1999; Taguchi and Kinzie, 2001), even though nitrate is usually more abundant in the water overlying reefs (Marubini and Davies, 1996). However, most studies reported that increased DIN can increase Symbiodiniaceae density, increase the contents of N and chrolophyll *a* per Symbiodiniaceae cell (Snidvongs et al., 1994), while elevated DIP did not affect Symbiodiniaceae densities (Muscatine et al., 1989), suggesting that Symbiodiniacea are sometimes in a state of DIN deficiency within the coral host. Yet there is relatively rare documentation in the literature about how Symbiodiniaceae species respond to N-deficient condition and switch between nitrate and ammonium.

*Fugacium kawagutii* (formerly *Symbiodinium kawagutii*, Clade F) is one of the Symbiodiniaceae dinoflagellates and was originally collected from the scleractinian coral *Montipora verrucosa* in Hawaii (Trench et al., 1987). It is relatively thermal-resistant, able to hold stable growth rate and maximum quantum yield of PSII (*Fv/Fm*) under higher temperature than species in genera *Breviolum, Cladocopium* and *Effrenium* that have been examined (Krueger et al., 2014). In addition, this species possesses highly duplicated nutrient transporters including 13 phosphate transporter genes, 62 nitrate transporter genes, and 84 ammonium transporter genes than other species (Li et al., 2020a; updated genome data also available at http://sampgr.org.cn/index.php/search). Therefore, these physiological characteristics may enable *F. kawagutii* to thrive under future global climate and potentially decreasing nutrient availability in the ocean. Studies have been conducted on molecular response in *F. kawagutii* to environment stress, e.g. thermal stress, phosphorus deficiency, and trace metals deficiency (Lin et al., 2019; Li et al., 2020b), but no research has focused on N nutrients.

In this study, we conducted physiological, cytometric and transcriptomic analyses on *F. kawagutii* treated with six N conditions. The first objective was to identify differentially expressed genes (DEGs) that respond to N deficiency and interrogate N-dependent metabolic processes. The second objective was to characterize physiological and transcriptomic differences between nitrate-based and ammonium-based *F. kawagutii* populations. These results will provide insights into the response of N-dependent metabolic processes in *F. kawagutii* to N nutrient variations.

## 2. Materials and methods

### 2.1 Alga cultures and nitrogen treatments

*F. kawagutii* (strain CCMP2468, Clade F) originally isolated from the scleractinian coral *Montipora verrucosa* in Hawaii. The culture was grown in L1-medium (Guillard et al., 1993) containing antibiotic cocktail comparing 100 mg/L streptomycin, 100 mg/L kanamycin and 200 mg/L ampicillin. The medium was prepared using surface seawater collected from northern South China Sea, which was 0.22-µm filtered and autoclaved. Temperature was controlled at 25 °C and illumination was provided at 200 µE · m ^−2^ s ^−1^ under a 14h: 10h light: dark cycle. Prior to experiments, the culture was inoculated into N-deprived medium to induce nitrogen starvation. Experimental cultures were set up in 1 L culture flasks for N-depleted, nitrate-replete, ammonium-replete and mixture of ammonium and nitrate at 2:1, 1:1, and 1:2 molar ratios but all with the same total nitrogen concentration as in N-replete condition. Each condition was set up in triplicate. While the N-depleted culture received 10 µM nitrate, the other five conditions all received 176 µM nitrogen. Cell counts were monitored daily using a Sedgwick-Rafter counting chamber (Phycotech, St. Joseph, MI, United States). Nitrate and ammonium concentrations in each culture were also determined daily by filtering 20mL culture through a 0.22 µm glass microfiber filters (GF/F) and subjecting the filtrate to measurements using cadmium column reduction photometric and indophenol blue photometric, respectively.

### 2.2 Measurement of chlorophyll a and photochemical efficiency (Fv/Fm ratio)

From each culture, a 20mL sample was filtered onto a 25mm GF/F filter and stored in a 5mL tube. Then the filter was immersed in 93% acetone and kept at 4 °C in the dark for about 48h to extract Chlorophyll *a*. Next, Chlorophyll *a* in the supernatant was measured using Turner Trilogy (Turner Designs fluorometer, USA) and averaged to per cell content. In addition, a 2mL sample was collected from each culture and kept in darkness for 20min, after which photochemical efficiency was measured using FiRe Fluorometer System (Satlantic LP, Halifax, Nova Scotia, Canada).

### 2.3 Measurement of cellular carbon and nitrogen contents

Samples were collected on day 5 and day 7 from mixture of ammonium and nitrate at 1:1 molar ratios group for cellular carbon (C) and N contents following previously protocol (Ducklow and Dickson, 1994). Briefly, 20mL sample was filtered onto a 25mm GF/F filter which had been combusted at 450 °C for 5h in a Muffle Furnace. Then the cell-containing filter was dried in 50 °C oven for over 12h and subjected to acid fumigation overnight at room temperature. Lastly, after the cell-containing filter was dried again and combusted, C and N were measured on vario PYRO-isoprime100 (Elementar, Germany), and the weight of each element was averaged to per cell content.

### 2.4 RNA extraction

Samples were collected on day 5 and day 7 from mixture of ammonium and nitrate at 1:1 molar ratios group and N-deficient group for RNA-seq. More than 3 × 10^6^ cells were harvested for each sample and total RNA was isolated previously reported (Li, et al., 2018, Li et al., 2020b). Briefly, after all cells were completely broken using with bead-beating (0.5mm mixed 0.1 mm diameter ceramic beads at 3:1) on the Fastprep^®^-24 Sample Preparation System (MP Biomedicals, USA), run for 3 cycles each at the rate of 6M/s for 1min, total RNA was extracted using Trizol regent (Molecular Research Center, Inc., USA) coupled with further purification using DiRect-Zol RNA miniprep (Zymo Research, Orange, CA, USA). RNA concentration was measured using a NanoDrop (ND-2000 spectrophotometer; Thermo Scientific, Wilmington, DE, USA), while integrity was assessed using an Agilent 2100 Bioanalyzer (Agilent Technologies, Palo Alto, CA, USA).

### 2.5 Illumina RNA-Seq and DEGs analysis

Samples with the RNA integrity number (RIN) ≥ 7 were used for RNA sequencing. One µg total RNA from each sample was used to generate a paired-end RNA-seq library by polyA mRNA workflow using TruSeqTM RNA sample preparation Kit from Illumina (San Diego, CA). Each library was subjected to sequence in Illumina Novaseq 6000 (2 × 150bp read length) at Majorbio Bio-pharm Technology Co., Ltd (Shanghai, China) with a final data out of 6Gbp for each sample. Clean data was obtained after removing low quality reads, reads containing only adapter, and reads containing poly-N in all the experimental samples. All the raw data from this study was deposited in the GenBank’s Sequence Read Archive (SRA) database (http://www.ncbi.nlm.nih.gov/) under the accession number xxxxxxx.

The latest version of the genome and gene set of *F. kawagutii* (Li et al., 2020a) were used as the reference database, which were available in the website of Symbiodiniaceae and Algal Genomic Resource database (SAGER, http://sampgr.org.cn) (Yu et al., 2020, unpublished data). The clean data were mapped to the genome and its updated gene set Fugka_Geneset_V3 using Hisat 2 (Perter et al., 2016) and RESM (Li and Dewey, 2011). Gene expression levels were calculated for each sample and genes with fold change ≥ 2 and P adjust value < 0.05 were accepted as differentially expressed genes (DEGs). Functional enrichment analyses including GO and KEGG were carried out using Goatools (https://github.com/tanghaibao/Goatools) and KOBAS 2.1.1 (Xie et al., 2011).

### 2.6 Core gene set update

Stably and highly expressed genes across three N treatment conditions were identified as core genes, following previously reported methods (Lin et al., 2019; Li et al., 2020a). Briefly, genes showing expression of ≥ 90% average Transcripts Per Million (TPM) and coefficient of variation (CV) ≤ 0.08 across all nitrogen treatment conditions were collected. Then, these core genes were compared with the previous core gene set version 2 (Li et al., 2020a) to generate core gene set version 3. To further rank the stability of expression level among top ten core genes, for potential “housekeeping” genes for future gene expression studies, log_2_ (TPM) were used to calculate the average expression stability value (M) by using geNorm (Vandesompele et al., 2002).

### 2.7 Statistical analysis

To evaluate the statistical significance of the differences observed between different nitrogen treatment groups, analysis of variance was performed using SPSS and Prism 8 statistics software packages. All data presented in this study are means with standard deviation calculated from the triplicated cultures in each nitrogen treatment group.

## 3. Result

### 3.1 Effect of nitrogen on F. kawagutii growth

Almost the same growth pattern was observed for NO_3_^−^ (NO3) and the NH_4_^+^ (NH4) treatment groups with maximum cell concentration at 148,737 and 158,420 cells mL^−1^ respectively in the 8-day experimental periods, except that there appeared to be a one-day lag for the NH4 treatment group (Figure 1A). While in the nitrogen-depleted group, cell concentration reached a plateau in 4 days with the much lower maximum concentration at 76,323 cells mL^−1^ during the experimental period (Figure 1A). Culture groups with mixture of ammonium and nitrate at 2:1, 1:1, and 1:2 molar ratios showed a similar growth curves with exponential growth from day 4 to day 8 reaching maximum cell concentrations on day 8 at 194,251, 204,010 and 198,385 cells mL^−1^ respectively (Figure 1B).

**Figure 1.**
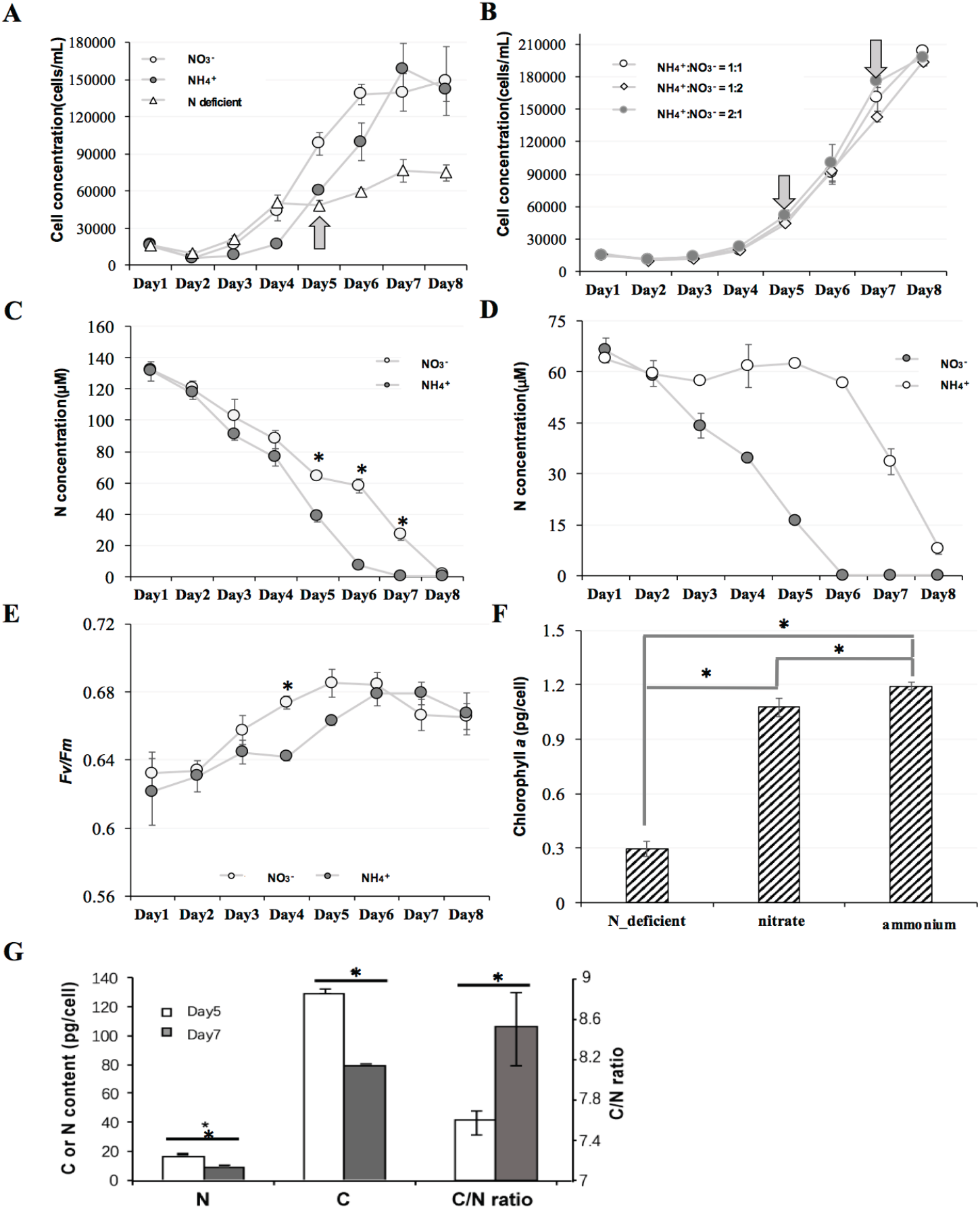
Growth cures (A,B), concentrations of remaining NH_4_^+^ and NO_3_^−^ (C,D), *Fv/Fm* (E), cellular Chlorophyll *a* content (F), as well as cell carbon, nitrogen contents and C:N ratio (G) of *Fugacium kawagutii*. **C** and **E** from NO_3_^−^ and NH_4_^+^ treatment group, while **D, F** and **G** from mixture of ammonium and nitrate 1:1 molar ratio group and N-deficient treatment group. Data points are means and error bars depict standard deviations of the triplicated cultures. Arrows indicate timing when samples were collected for RNA-seq and asterisks depict significant differences between the two comparison groups.

### 3.2 Differential ammonium and nitrate uptake and photosynthetic parameters

In the NO3 and NH4 groups, although staring from the similar initial concentration of nitrate and ammonium, they started to differ on the fifth day (P < 0.05, two-way ANOVA) and the cells took up ammonium faster than nitrate (P < 0.05, two-way ANOVA, Figure 1C). When both ammonium and nitrate were present at 1:1 ratio (NO3NH4) and at equivalent total N concentration to the NO3 or NH4 group, cells preferentially absorbed ammonium as its concentration remaining in the medium decreased rapidly whereas the concentration of remaining nitrate in the culture medium stayed almost unchanged until day 5, when remaining NH_4_^+^ dropped below detection limit (Figure 1D). After day 5, cells began to absorb nitrate as its concentration remaining in the medium decreased rapidly over the next three days. This mode of uptake also occurred in other NO3NH4 groups in which the ammonium to nitrate ratio was 1:2 or 2:1 (Figure S1).

Furthermore, the NO3 and NH4 groups exhibited generally similar maximum quantum efficiency (*Fv/Fm*) although a lower efficiency was observed in the NH_4_ group on one day (P < 0.05, two-way ANOVA) for unknown reason (Figure, 1E). In the NO3NH4 cultures, day 5 was when the cells only absorbed ammonium, while day 7 was when the cells only absorbed nitrate (Figure 1D). Cellular content of chlorophyll *a* was significantly lower in the N-depleted group than the N-replete groups (P < 0.05, t-test) and was higher on day 5 (NH_4_^+^) than day 7 (NO_3_^−^) (P < 0.05, t-test) (Figure 1F). In addition, cell contents of carbon and nitrogen were higher on day 5 than on day 7, and so was the C: N ratio (P < 0.05, t-test, Figure 1G).

### 3.3 Transcriptomic response to nitrogen deficiency

Samples were collected for RNA-seq from the NO3NH4 group on day 5 (only ammonium was utilized) and day 7 (only nitrate was utilized), and the N-depleted group (Figure 1). RNA-seq yielded average 54,369,734 clean reads from each of the three triplicated (totally 9) cultures (Table S1). Approximately 82.88% of the RNA-seq reads had matches in the recently improved genome of *F. kawagutii* (Table S1). Principal component analysis (PCA) for the transcriptomic data indicated that the within-group biological replicates were close to each other whereas different treatment groups were well separated from each other, validating the experimental setup and reliability of the RNA-seq data for differentially expressed genes (DEGs) analysis (Figure 2A). The RNA-seq data yielded 45,192 genes. A number of genes were highly expressed across all three treatments, including ferredoxin, peridinin chlorophyll-a binding protein apoprotein precursor (PCP), nitrate transporter, cytochrome c-550, phosphoglycerate kinase, ribulose bisphosphate carboxylase/oxygenase (rbcL), pre-mRNA-processing factor 6, and major basic nuclear protein 2 (Table S2). A total of 3,230 genes were differentially expressed between the N treatment conditions with a cutoff of log_2_(fold change) > 1 and p-value adjust < 0.05. Similar number of DEGs were found for NO_3_^−^ vs N-deficient (2,515) and NH_4_^+^ vs. N-deficient (2,507) comparisons, but only a very small number of DEGs (33) were detected for the NO_3_^−^ vs. NH_4_^+^ comparison (Figure 2B, C and D). There were more down-regulated genes than up-regulated genes across all three treatment comparisons (Figure 2B and D).

**Figure 2.**
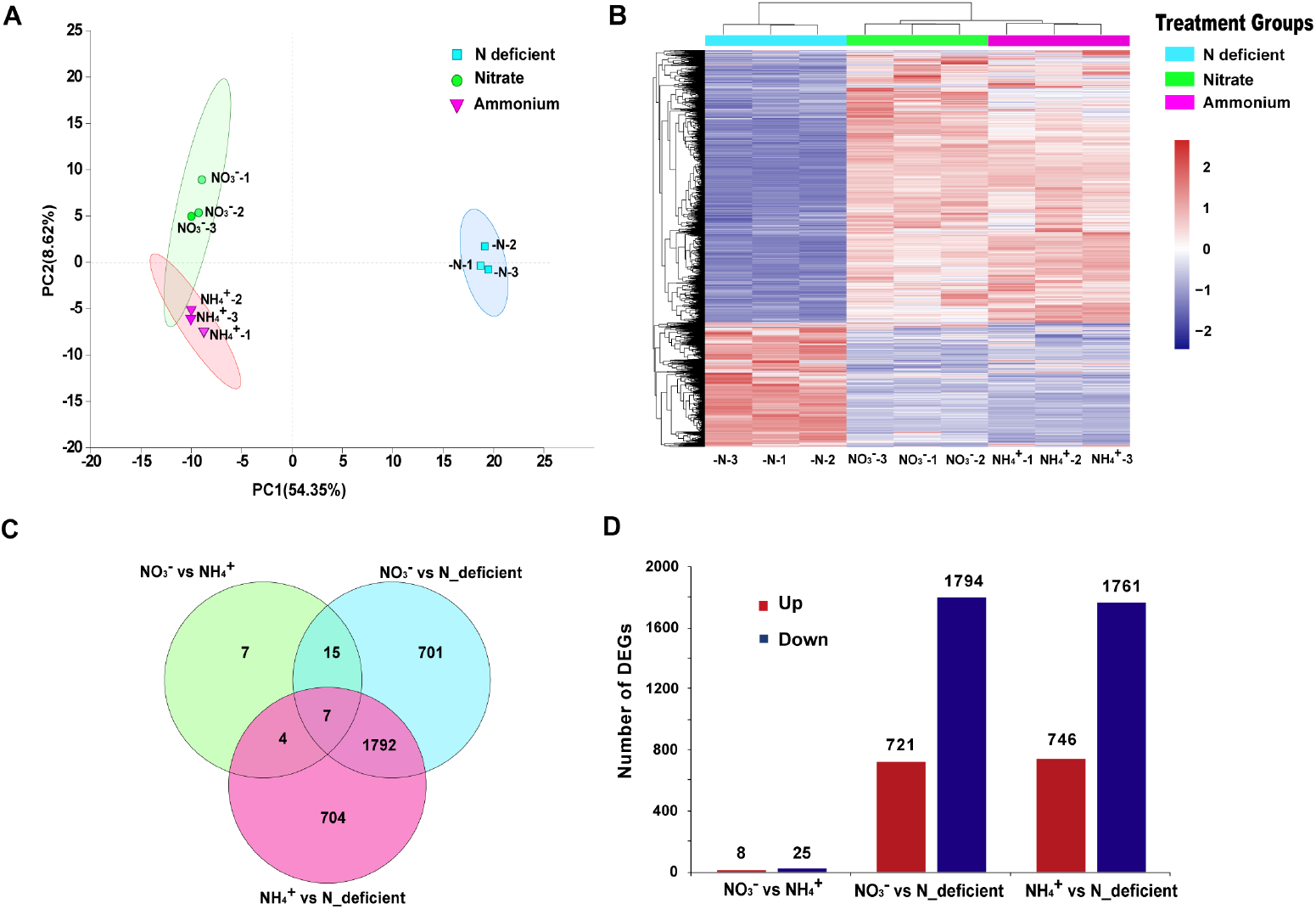
Distinct transcriptomic profiles of *F. kawagutii* under three different N-nutrient conditions. **(A)** Principal component analysis of the RNA-Seq data of *F. kawagutii* showing separation of the three treatments with small differences among within-group replicates (labeled with -1, -2, -3). **(B)** Heatmap of all differentially expressed genes (DEGs) from three treatment comparisons. **(C)** Venn diagram showing shared and unique DEGs among the three treatment comparisons. **(D)** Histogram showing the number of up- and down-regulated DEGs in each comparison group.

Because only 33 DEGs came from the NO_3_^−^ vs. NH_4_^+^ comparison, there was no significantly enriched GO terms and KEGG pathway found from this comparison. The DEGs from N-replete vs. N depleted comparison were significantly enriched in various GO terms (Figure 3A, Table S3), indicating a broad metabolic impact by N-nutrient variabilities. The most remarkable Biological Processes enriched by the DEGs were nitrogen cycle metabolic process and nitrate assimilation and carbon fixation. For Cellular Component, chloroplast, plastid, and ribosomal subunit were most highly enriched by DEGs. At the Molecular function level, glutamate synthase (NADH) activity, phospholipid binding and ammonium transmembrane transporter activity were highly represented in the DEGs. KEGG analysis, on the other hand, showed that ribosome, nitrogen metabolism, photosynthesis and mRNA surveillance pathway were significantly enriched (Figure 3B). From these enriched GO terms and KEGG pathways, several themes emerged.

**Figure 3.**
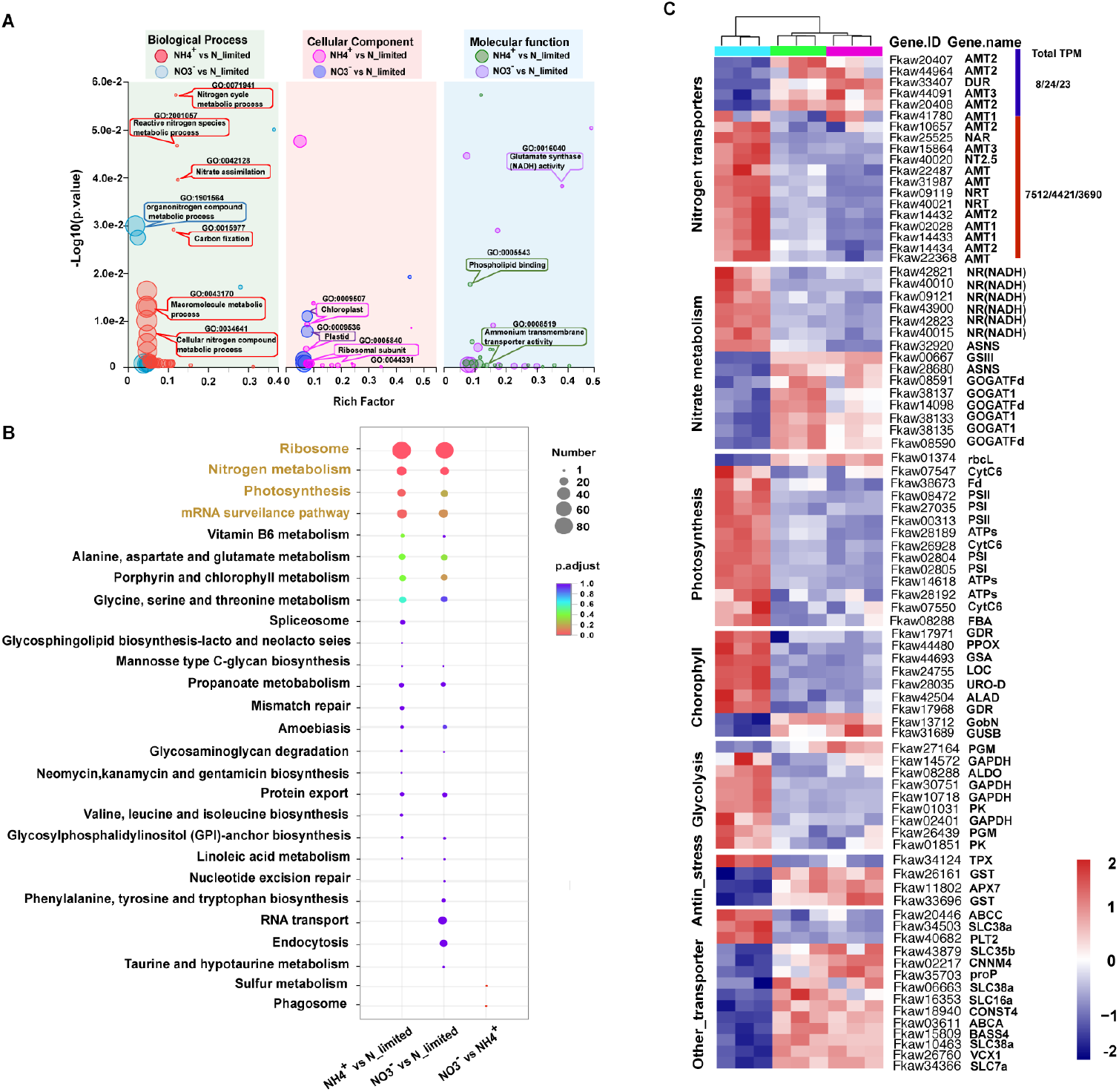
Select GO terms, KEGG pathways and key genes that responded to nitrogen deficiency in *F. kawagutii*. **A)** GO enrichment analysis of DEGs in each comparison group; significantly enriched GO terms (q-value < 0.05) are shown. Three term types are highlighted with green, red and blue respectively. Most interesting Go terms are shown in text boxes. The details of each GO term were summarized in Table S3. **B)** KEGG enrichment analysis of DEGs in each comparison group; significantly enriched KEGG pathways are in color fonts. **C)** Heatmaps illustrating the expression levels of select DEGs in all samples (top horizontal bars: cyan, N-deficient; green, NH_4_^+^; pink, NO_3_^−^) based on RNA-Seq. The heatmap color strength represents normalized gene expression (TPM), from blue (lowest), white, to red (highest). Total TPM on the upper right indicate the summed TPM values of the up-regulated genes (grouped by the red vertical bar on its nearest left) and down-regulated genes (grouped by the blue vertical bar on its nearest left) under N-deficient/NH4/NO3 conditions. The detail of these genes is summarized in Table S4.

The first emergent theme was about N-responsive metabolic pathways. Totally 52 genes involved in nitrogen transport and metabolism were found in the N-depleted group compared with N-replete group, which were defined as nitrogen responsive genes in *F. kawagutii* (Table S4). Among these 52 N responsive genes, sixteen of the twenty genes encoding ammonium transporters were up-regulated under N deficiency, whereas one ammonium transporter 3 gene and three ammonium transporter 2 genes were down-regulated. Thirteen nitrate transporter genes and three nitrite transporters genes were up-regulated under N deficiency. Only one urea symporter (*DUR*) gene was among the DEGs and it was down-regulated under N deficiency. In addition, eight other transporter genes, including sodium-coupled neutral amino acid transporter 6, were up-regulated, and sixteen others were down-regulated, including GDP-mannose transporter, cationic amino acid transporter 1, proline transporter, sodium/metabolite cotransporter and ABC transporter A family. Further analysis on intracellular transformation processes of nitrogen indicated that six nitrate reductase [NADH] genes and one asparagine synthetase (*ASNS*) gene were up-regulated, while three glutamate synthase 1 [NADH] genes (*GOGAT1*), three ferredoxin-dependent glutamate synthase genes (*GOGAT_Fd*), one glutamine synthetase (*GSIII*) gene, and another asparagine synthetase gene were down-regulated under N deficiency (Figure 3C, Table S4). These results indicated the reconfiguration of N-nutrient acquisition machinery to promote uptake of available nitrogen and to reduce investment in the assimilation apparatus due to the limited availability of nitrogen.

Nine of these 52 N-responsive genes were regulated under N-deficient condition but not under other eight treatment conditions including five trace metal deficiency (Li et al., 2020b), phosphorus deficiency and organic phosphorus replacement treatment conditions in *F. kawagutii* (Lin et al., 2019). They encode ammonium transporter1, nitrate transporter, nitrate reductase, nitrite transporter, and type-3 glutamine synthetase. These are specific to N-stress in *F. kawagutii* (Figure 4, Table S5).

**Figure 4.**
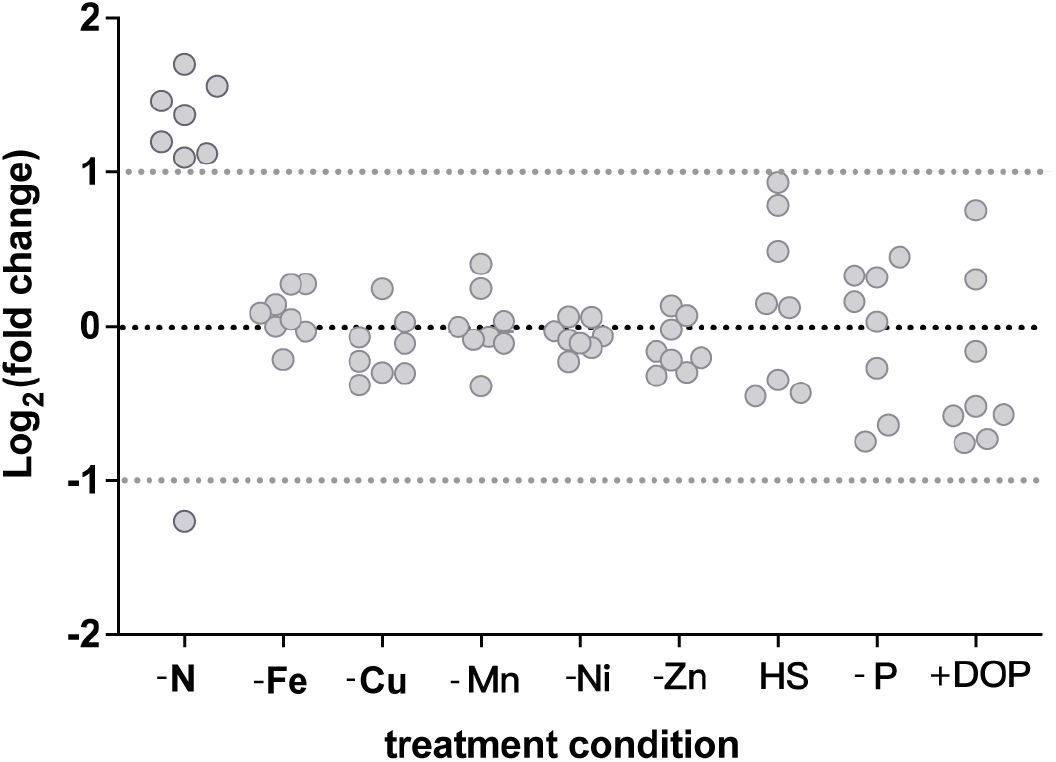
Expression fold change of N-stress specific genes in response to nine nutrient conditions. Horizontal line depicts no change in expression, above line are genes up-regulated and below line are genes down-regulated under nine treatment conditions. -N: nitrogen deficiency; -Fe: iron deficiency; -Cu: copper deficiency; -Mn: manganese deficiency; -Ni: nickel deficiency; -Zn: zinc deficiency; HS: heat stress; -P: phosphorus deficiency; +DOP: organic phosphorus addition. Detail of these nine genes were summarized in Table S5. Data from Lin et al., 2019 and Li et al., 2020b.

The second emergent theme was with photosynthesis. Most genes involved in the photosynthesis electron transfer chain (ETC) were up-regulated under N deficiency. These included eight genes encoding photosystem II protein, three genes encoding photosystem I protein, four genes encoding chloroplast ATP synthase, one gene encoding ferredoxin, one gene encoding cytochrome b6-f complex iron sulfur subunit and two gene encoding cytochrome c6 were all up-regulated (Figure 3C, Table S4). On the contrary, the expression of the form II Rubisco (rbcL), an essential gene for CO_2_ fixation in the Calvin cycle decreased under N deficiency. The strengthened electron transport system and weakened ability to fix CO_2_ might indicate the potential of homeostasis disruption of energy and reducing power in the cell. ETC is the main source of reactive oxygen species (ROS) production in photosynthetic organisms (Cleland and Grace, 1999), where the instability of electron transport may cause electrons to leak out and combine with oxygen to form ROS.

The third emergent theme lied in the cell cycle and anti-stress. Analysis of cell division related genes indicated that 10 genes was down-regulated in the N-depleted group, including cyclin-dependent kinase 10, dual specificity protein phosphatase 3, mitotic checkpoint protein and mitogen-activated protein kinase kinase (MAPKK) (Figure 3C, Table S4). This was consistent with the lower growth rate under N deficiency. Ribosome biogenesis and protein translation are finely coordinated with and essential for cell growth and proliferation. In this study, we found 7 genes encoding 30S ribosomal protein and 10 (out of 11) genes encoding 50S ribosomal protein were down-regulated under N deficiency, while 15 (out of 19) genes encoding 40S ribosomal protein and 36 (out of 39) genes encoding 60S protein were up-regulated under N deficiency (Figure 5A, Table S4). This was indicative of decreased organellar protein synthesis capacity while increasing cytoplasmic protein synthesis capacity as a response to N deficiency. Analysis of ROS scavenging genes found three genes encoding L-ascorbate peroxidase 7, glutathione S-transferase and glutathione S-transferase class-mu were down-regulated and only one gene encoding thioredoxin was up-regulated under N deficiency (Figure 3C, Table S4), indicating a potential that ROS scavenging mechanism was altered and the capacity might be lowered under N deficiency.

**Figure 5.**
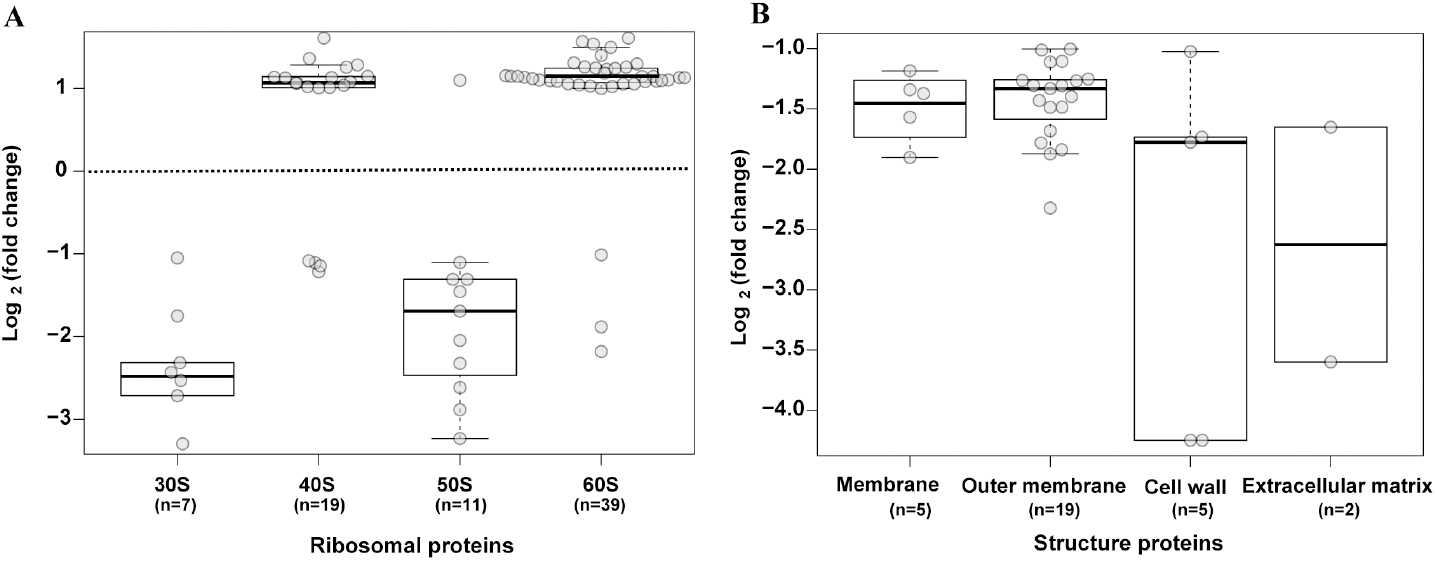
Log transformed fold change of the genes encoding ribosomal proteins (A) and cell surface structure proteins (B) under N deficient condition in *F. kawagutii*. Numbers in parentheses (n) represent the numbers of DEGs in that category. The dots represent the fold change of every gene.

The fourth emergent theme had to do with glycolysis and tricarboxylic cycle (TCA). For glycolysis, one gene encoding phosphoglycerate mutase was down-regulated, while another phosphoglycerate mutase gene, along with five glyceraldehyde-3-phosphate dehydrogenase genes (GAPDH), one fructose-bisphosphate aldolase genes and two pyruvate kinase genes were up-regulated under N deficiency (Figure 3C, Table S4). Among these genes, pyruvate kinase is one of the rate-limiting enzymes in glycolysis, and the other two rate-limiting enzymes, hexokinase and phosphofructokinase, displayed no significant change in transcript abundance between the N conditions. These together signified an uptick of glycolysis under nitrogen deficiency. For TCA analysis, only citrate synthase gene exhibited a moderate down-regulation (fold change between 1.5 and 2, P < 0.05) under nitrogen deficiency, whereas other two key enzymes, isocitrate dehydrogenase and α-ketoglutarate dehydrogenase, showed no differential expression. Apparently, TCA was down tuned under N deficiency.

The fifth emergent theme is with the cell wall and cell membrane. From our transcriptomic data, five genes encoding membrane proteins, nineteen genes encoding outer membrane proteins, five genes encoding cell wall protein and two genes encoding extracellular matrix proteins were all down-regulated under N deficiency. These were consistent evidence that cell surface structure was affected by N deficiency (Figure 5B, Table S4).

### 3.4 Transcriptomic response to the switch from ammonium to nitrate utilization

From our transcriptomic data, 256 genes showed strong (33; fold change ≥ 2, P < 0.05, as defined earlier in the paper) or moderate regulation (223; fold change between 1.5 and 2, P < 0.05) under ammonium condition compared with nitrate condition (Table S6). Of these, molecular functions of ammonium transport (four ammonium transporter 1 genes) and assimilation (two glutamate synthase 1 [NADH] gene, and one ferredoxin-dependent glutamate synthase gene) were down-regulated in the NH_4_^+^ group (Figure 6A).

**Figure 6.**
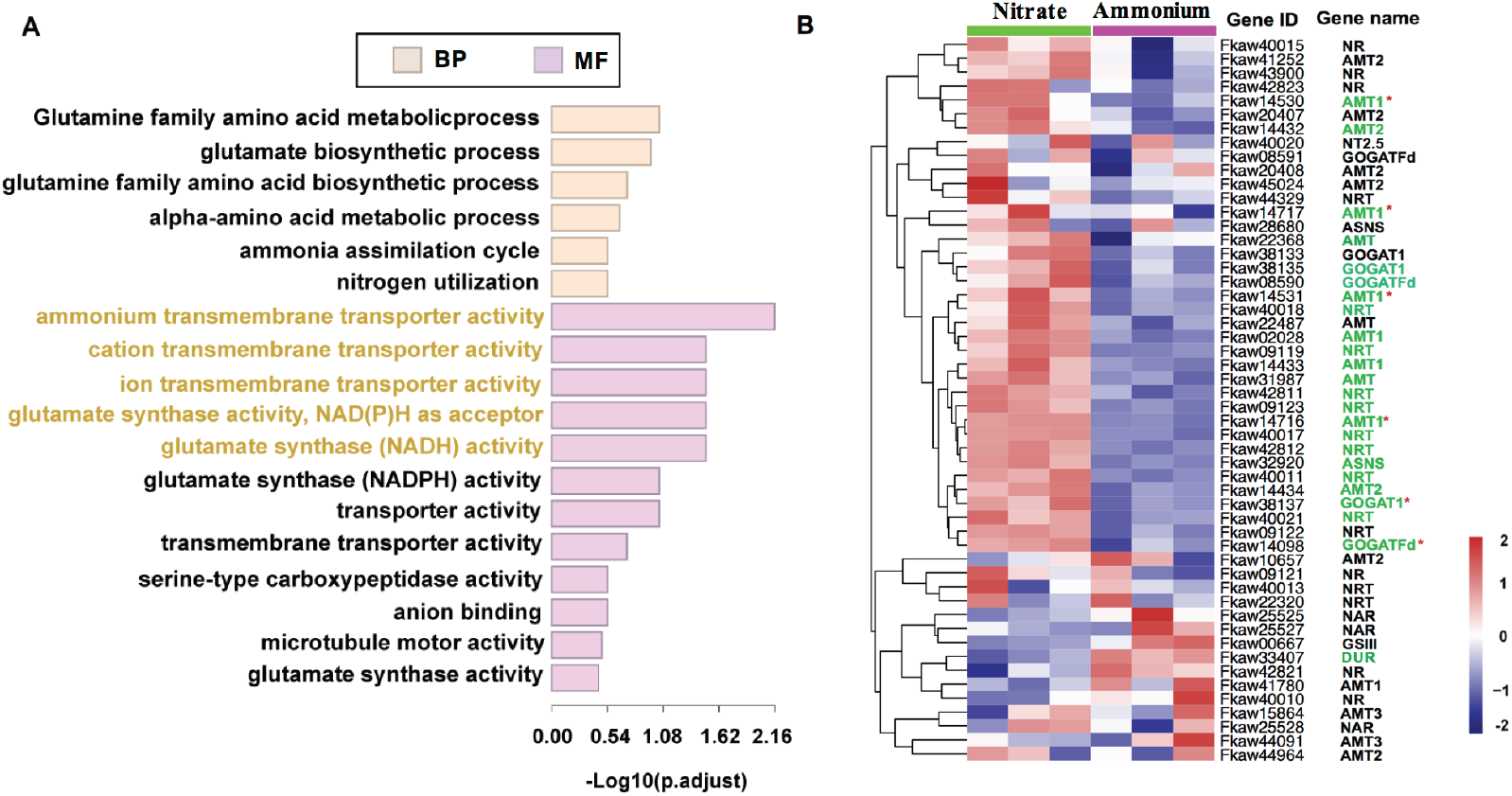
Several important DEGs-enriched GO terms and the expression of 52 nitrogen responsive genes under ammonium utilization relative to nitrate utilization in *F. kawagutii*. **A)** GO enrichment of DEGs (fold change > 1.5, P < 0.05). Significantly enriched GO terms are in color fonts. **B)** Heatmap illustrating the expression of 52 nitrogen responsive genes under nitrate and ammonium conditions. Genes showing fold change > 1.2 and P < 0.05 are in green fonts and those above 1.5 are also marked with an asterisk. NR: nitrate reductase; AMT: ammonium transporter; NT2.5: high affinity nitrate transporter; GOGAT_Fd: ferredoxin-dependent glutamate synthase; NRT: nitrate transporter; ASNS: asparagine synthetase; GOGAT1: glutamate synthase 1[NADH]; *NAR*: nitrite transporter; GSIII: type-3 glutamine synthetase; DUR: urea symporter.

From the 52 N uptake and assimilation related genes that were identified as responsive to N-deficient condition, 24 of them were also responsive to the switch from ammonium to nitrate conditions (Figure 6B, Table S7), but the fold changes of these genes were small (below 2). Among these 24 genes, 6 gene showed fold changes between 1.5 and 2 and 18 genes showed fold changes between 1.2 and 1.5 (P < 0.05) (Figure 6B, Table S7). Only DUR was up-regulated and the other 23 genes were down-regulated under ammonium condition (Figure 6B), indicating nitrogen transport and metabolism were down-regulated when ammonium was the major N nutrient. In addition, analysis of ROS scavenging genes revealed that genes encoding ascorbate peroxidase (fold change between 1.5 and 2, P < 0.05), glutathione S-transferase and peroxiredoxin (fold changes between 1.2 and 1.5, P < 0.05) were moderately up-regulated under NH_4_^+^ growth condition.

### 3.5 Core gene set update

We found a total of 1,176 genes that exhibited stable and high expression levels across the three N conditions based on our strict criteria (Table S8). The expression of these core genes averaged 66.38 TPM and ranged from 35.12 TPM to 1005.6 TPM. Among these, 1,067 (90.73%) genes were functionally annotatable. As an effort to develop a core gene repertoire for this species, we have previously identified 115 common core genes, which were highly and commonly expressed under nine different growth conditions (temperature, phosphate, and trace metals), which was denoted as core gene set version 2 (Li et al., 2020a). Here, we combined core gene set version 2 with present core genes, which yielded 10 common core genes, now across 11 conditions, herein denoted as core gene set version 3 (Figure 7A, Table S9). Among these ten genes, tubulin was most highly expressed and E3 ubiquitin-protein ligase was least highly expressed (Figure 7A). Based on the expression stability analysis using geNorm, all the ten core genes showed far smaller variability (M value) than the threshold value 1.5, indicating that the expression levels of these genes were relatively stable under three nitrogen deficiencies (Figure 7B). Ranked as the most stable genes were tyrosyl-DNA phosphodiesterase 2 and light-harvesting complex I.

**Figure 7.**
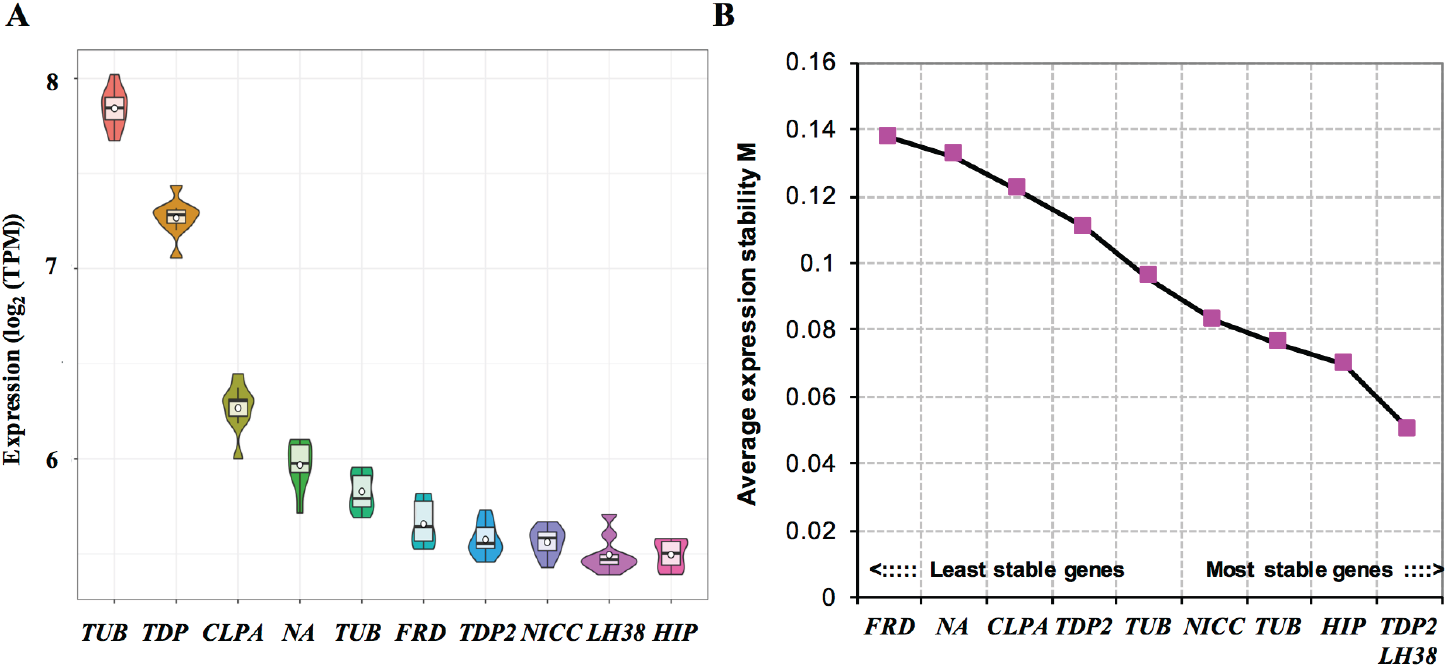
Expression level and stability of the ten core genes under three nitrogen conditions. **A**) Boxes plot of gene expression levels (TPM with log_2_ transformation). **B**) The stability and its ranking of gene expression calculated using geNorm based on expression shown in **A**. TUB: tubulin; TDP2: Tyrosyl-DNA phosphodiesterase 2; CLPA: ATP-dependent Clp protease ATP-binding subunit; NA: not annotated; FRD: fumarate reductase; NICC: 6-hydroxynicotinate 3-monooxygenase; LH38: light-harvesting complex I; HIP: E3 ubiquitin-protein ligase.

## 4. Discussion

Nitrogen (N) as a crucial component of amino acids, proteins, nucleotides (ATP, cAMP), nucleic acids and chlorophylls is a key macronutrient for photosynthetic productivity in the ocean. In the symbiotic system, the coral host can directly provide its metabolic waste of N to the endosymbionts (Davy et al., 2012). Therefore, symbiotic Symbiodiniaceae N nutrient supply is not only subjected to variability in the ambient environment but also to coral’s physiological fluctuation. Understanding how Symbiodiniaceae respond to the dynamics of N abundance and stoichiometric proportions of ammonium and nitrate is critical to assessing how the health status of the symbiosis is influenced by the natural environment and the host physiology. By investigating physiological, cytometric, and transcriptomic responses in *F. kawagutii* to N-deficiency and switch between ammonium and nitrate, in this study we mainly found that N-deficiency caused growth depression, lower energy production, and less investment in photosynthate transport, and growing on ammonium produced a similar cell yield as nitrate, but with a decreased investment in nutrient transport and assimilation. These all have important implications to energy and photosynthate translocation to support symbiosis. Besides, by integrating our current results with previous data, we identified nine candidate N-deficiency marker genes which were only regulated under N-deficient condition, and ten highly and stably expressed genes as candidate reference genes that will be useful for future gene expression studies.

### 4.1 Depressed growth, photosynthesis, antioxidant capacity, and symbiosis potential under N deficiency

Decreased cell growth, cellular chlorophyll *a* and cellular N content are typical responses to N deficiency in most phytoplankton e.g. *Dunaliella tertiolecta* (Berges and Falkowski, 1998; Geider et al., 1998), *Thalassiosira pseudonana* (Chen et al., 2018), *Porphyridium cruentum* (Zhao et al., 2017), and Symbiodiniaceae (Radecker et al., 2015). Therefore, the lower growth rate, cellular Chl *a* content observed in the N-depleted cultures of *F. kawagutii* in the present study were as expected. Under N deficiency, a set of cell division genes was down-regulated, consistent with slower growth rate, whereas most DEGs involved in Chorophyll synthesis and electron transfer function were up-regulated, while cellular Chl *a* content decreased, indicating the cells’ response by promoting energy and reductant production under N deficiency. In addition, ribosome biogenesis and protein translation are finely coordinated with and essential for cell proliferation and differentiation (Zhou et al., 2015). Our transcriptomic data indicate most genes encoding 30S and 50S ribosomal proteins were down-regulated under N deficiency, consistent with lowered growth rate, but in contrast with the pattern of elevated 40S and 60S ribosomal protein expression. The increased expression of the 40S and 60S ribosomal proteins may be a response to the needs for increasing synthesis of N transporters and assimilation related enzymes. This is true even for most transporters and enzymes functioning in the organelles, because for typical dinoflagellates such as Symbiodiniaceae most of the mitochondrial and chloroplast proteins are encoded in the nucleus and synthesized with the 40S/60S ribosomes (Lin, 2011).

As rbcL gene was significantly down-regulated, photosynthetic carbon fixation was likely reduced under N deficiency. However, most DEGs involved in glycolysis were up-regulated under N deficiency, e.g. pyruvate kinase, phosphoglycerate mutase, fructose-bisphosphate aldolase, GAPDH, suggesting that the N-limited cells need to consume more glucose to cope with N deficiency. The reduction of photosynthate production and the increase in consumption of glucose in Symbiodiniaceae would have significant implications to its symbiosis with corals or other hosts, as it may result in a reduction in the output of photosynthates to the host. Notably, cationic amino acid transporter and proline/betaine transporter genes were also down-regulated. Together, these results suggest that *F. kawagutii* may be more parasitic than symbiotic under N deficiency.

Reactive oxygen species (ROS) have been suggested as a culprit of thermal stress and solar radiation-induced bleaching (Lesser, 1997). ROS is mainly formed by the inevitable leakage of electrons onto O_2_ from the ETC of chloroplast and mitochondria in plants (Sharma et al., 2012). As long as scavenging mechanisms are functional, ROS will be removed by antioxidant systems (Marty-Rivera et al., 2018). Our transcriptomic data indicated that DEGs encoding L-ascorbate peroxidase 7, glutathione S-transferase were down-regulated under nitrogen deficiency. *F. kawagutii* contains a diverse set of SOD (Cu/Zn, Mn/Fe, and Ni-dependent) encoding genes (Lin et al., 2015), but none of them showed differential expression between N-depleted and N-replete conditions. These results combined suggest that antioxidant capacity decreased under N deficiency. Our previous study also found glutathione S-transferase and ascorbate peroxidase were down-regulated under iron deficiency (Li et al., 2020b), indicating that iron and nitrogen are both important in resisting oxidative damage.

For Symbiodiniaceae as endosymbionts, cell surface structure is very important to maintain symbiotic relationship (Davy et al., 2012; Xiang et al., 2015; Xiang et al., 2017). Our previous studies found most of the genes identified encoding cell surface structure proteins were regulated under deficiency of iron and other trace metals (Li et al., 2020b). In the present study, genes associated with cell wall proteins, extracellular matrix proteins and cell membrane proteins were down-regulated under N deficiency, reveling that the cell surface structure is shaped by N-nutrient availability. This, along with the reduced antioxidative capacity discussed above and previous findings suggest that nutrient deficiency can lead to changes in the ability of ROS scavenging and cell surface structure of *F. kawagutii*, which may contribute to the breakdown of the relationship between coral and the Symbiodiniaceae alga.

### 4.2 N-deficiency responsive genes

In present study, 52 genes involved in nitrogen transporting and metabolism were regulated under N-deficient condition, which were regarded as N responsive gene in *F. kawagutii*. Among these genes, most genes encoding NRT and AMT were up-regulated under N deficiency, which is consistent with these roles in nitrogen uptake and assimilation. Comparison with previous transcriptomic data created under eight other different conditions, current data showed that 9 of the 52 genes, AMT included, appeared to exclusively respond to N deficiency. In contrast, genes encoding GOGAT_Fd, GOGAT1, GSIII and ASNS were down-regulated under N-deficient condition. All these results indicate the reconfiguration of N-nutrient acquisition machinery to promote uptake of available nitrogen and to reduce investment in the assimilation apparatus due to the limited availability of nitrogen. Regulation of *NRT* and *AMT* under nitrogen deficiency was found in most phytoplankton, e.g. *Karenia brevis* (Morey et al., 2011), *Prorocentrum shikokuense* (Li, unpublished data), *Thalassiosira pseudonana* (Chen et al., 2018). *Fragilariopsis cylindrus* (Bender et al., 2014), and *Pseudo-nitzschia multiseries* (Bender et al., 2014), while those involved in nitrogen metabolism as well as carbon metabolism were exhibited different expression patterns among these species. Thus, it appears that the regulation of these pathways may differ somewhat across different phytoplankton phyla.

### 4.3 Preference for ammonium over nitrate and molecular mechanisms

Despite the clear evidence that acquiring ammonium from seawater into the coral-Symbiodiniaceae symbiotic system may be a key component of coral nutrition (Pernice et al., 2012), the reason why ammonium as a nitrogen source is advantageous poorly understood for the symbiotic system. In this study, we found that *F. kawagutii* assimilated ammonium faster than nitrate when they were provided in separate cultures (Figure 1C). There are two possible explanations. Firstly, the number of gene encoding ammonium transporters (84) was higher than gene encoding nitrate transporters (62) in *F. kawagutii* (Table S10). Secondly, ammonium can also move across the plasma membrane through non-specific systems, including potassium channels/transporters (ten Hoopen, et al., 2010). The faster uptake of ammonium than nitrate resulted in the lower C: N ratio per cell in the ammonium-grown than the nitrate-grown cells (Figure 1G). Interesting, we found that not only ammonium was preferentially utilized but that nitrate uptake was significantly inhibited when both ammonium and nitrate were provided in the culture medium (Figure 1D). Similar findings have been reported on other phytoplankton (Glibert et al., 2016). In barley roots, the preferential uptake of ammonium was thought to involve high-affinity nitrate transport system that might be sensitive to ammonium (Kronzucker et al., 1999; Hachiya and Sakakibara, 2017). It has been shown that nitrate transporter in the land plant model *Arabidopsis thaliana* (*AtNRT 2.1*) is the target for ammonium-dependent inhibition of nitrate uptake (Cerezo et al., 2001). It is unclear if this mechanism exists in *F. kawagutii*, but our transcriptome data showed that most NRT encoding genes were down-regulated in ammonium-based cultures. In addition, although *F. kawagutii* preferentially absorbed ammonium and stocked more nitrogen (likely as amino acids or proteins) in the cell, higher concentrations of ammonium exhibited inhibitory effects on growth (Figure 1A). It is thus apparent that *F. kawagutii* cells absorbed and stored more ammonium under high concentration of ammonium at the expense of growth (Figure 1A, G).

Expression analysis indicate that 23 genes among the 52 N-stress responsive genes responsive to the switch from ammonium to nitrate utilization were moderately down-regulated under ammonium condition (Figure 6). Compared to ammonium utilization, nitrate needs to be reduced first to nitrite and then to ammonium before it can be assimilated into amino acids, which requires ATP and reductants derived from photosynthesis (Falkowski and Stone, 1975). This, along with lower expression of N transport and metabolism genes, it seems that using ammonium as N nutrient saves energy. This freed up energy may be reallocated to other metabolic needs, such as combat oxidative and maintain symbiotic relationship with corals under ammonium treatment condition (Figure 8). In support of this, an ascorbate peroxidase, glutathione S-transferase and peroxiredoxin genes were moderately up-regulated (P < 0.05) under ammonium condition.

**Figure 8.**
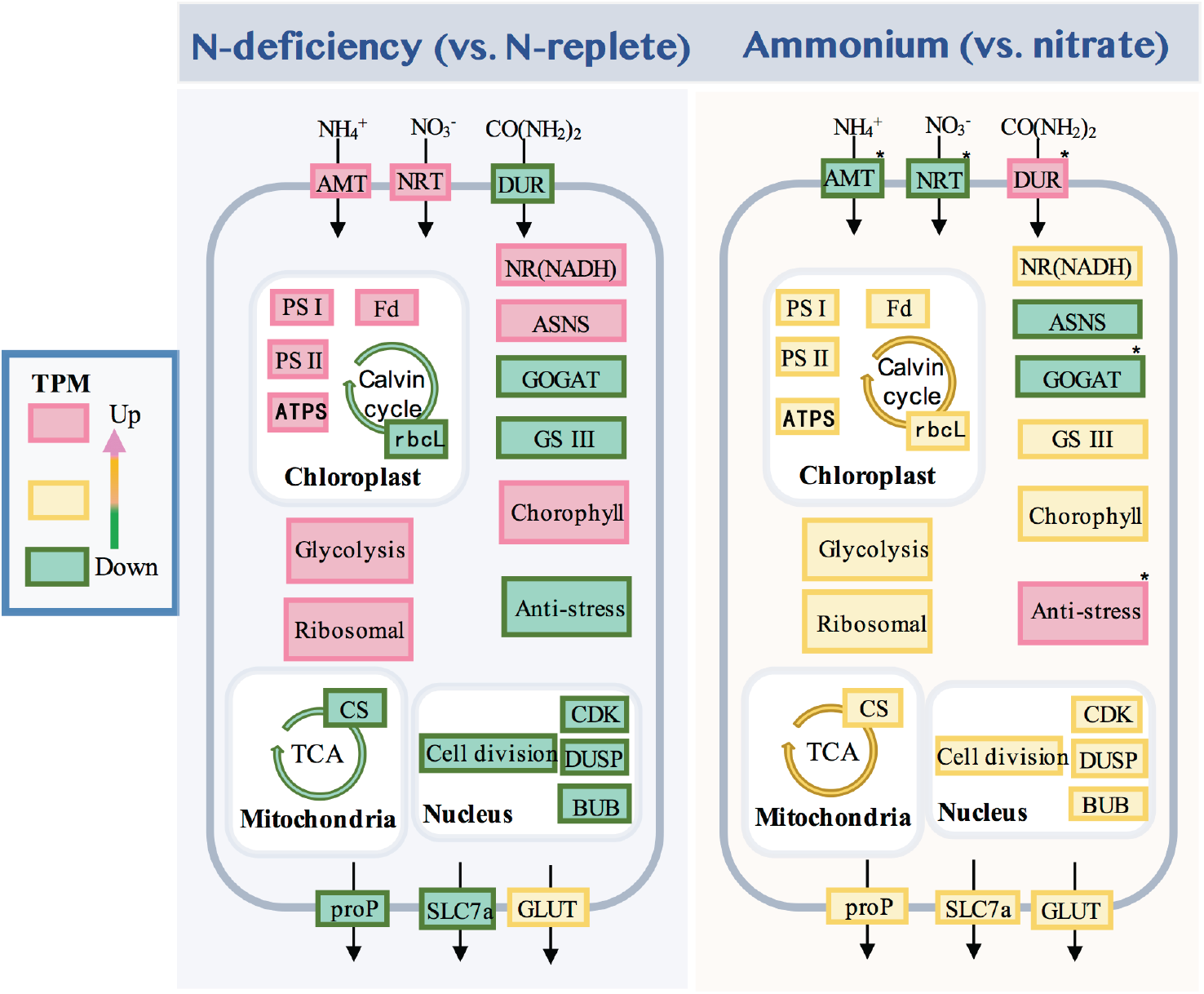
Schematic representation of metabolic pathways in *F. kawagutii* under nitrogen deficiency and ammonium utilized conditions. AMT: ammonium transporter; NRT: nitrate transporter; DUR, urea symporter; PSI: photosystem I protein; PSII: photosystem II protein; ATPS: ATP synthase; Fd: ferredoxin; rbcL: ribulose bisphosphate carboxylase; NR(NADH): nitrate reductase; GOGAT: glutamate synthase; MDH: malate dehydrogenase; CDK: cyclin-dependent kinase; DUSP: dual specificity protein phosphatase; BUB: mitotic checkpoint protein; proP: proline transporter; SLC7a, amino acid transporter; GLUT: glucose transporter; The asterisks represent genes with moderate regulation (fold change > 1.5, P < 0.05).

### 4.4 Highly expressed genes and core gene set update

A small number of genes are highly represented in the EST databases in dinoflagellates, e.g. *P. shikokuense* (Yu et al., 2019); *F. kawagutii* (Li et al., 2020b). In this study, we found ferredoxin, PCP, nitrate transporter, cytochrome c-550, phosphoglycerate kinase, rbcL, pre-mRNA-processing factor 6, and major basic nuclear protein 2 were highly expressed across all three nitrogen treatment conditions used in this study. Among these, PCP, nitrate transporter and rbcL were also highly expressed across five trace metal culture conditions (Li et al., 2020b), indicating that the *F. kawagutii* have a high ability of light harvesting and nutrient transporting. Besides, only < 30% of genes have been shown to exhibit transcriptional changes in response to environmental fluctuations or stress in dinoflagellates (Lin, 2011). Therefore, the small proportion of genes were differentially expressed under three N treatment conditions especially switch from ammonium to nitrate. In addition, we found while multiple unigenes encode the same functional protein, the highly expressed genes appeared to be stably expressed, but the genes with lower expression levels were clearly regulated in response to N nutrient condition changes (result not shown). This implicates that the cells respond to external pressure by regulating the low-expression genes in *F. kawagutii*. This is potentially interesting but requires further investigation.

By combining current data with previously reported data (Li et al., 2020a) and applying strict selection criteria, 10 genes were found to exhibit stable and high expression levels across 11 different conditions, including phosphorus nutrients, trace metals, and now nitrogen nutrients, which we designated as core gene set version 3. In this core gene set, there are two genes annotated as tubulin alpha chain and one of them showed highest expression level of all genes. Tubulin has been considered to be one of the most stable reference gene in animal (Zhai et al., 2014), plant (Zhang et al., 2018) and phytoplankton (Krueger et al., 2015). Therefore, the stable and high expression level under 11 treatment conditions makes tubulin a most suitable candidate reference gene for gene expression studies in *F. kawagutii*. In addition, GAPDH is another most commonly used reference genes (Kozera and Rapacz et al., 2013) and also validated by RT-qPCR in our previous study (Li et al., 2020a), but four genes encoding GAPDH were all up-regulated under N deficiency, making it not a good reference gene in *F. kawagutii* for nitrogen deficiency gene expression studies.

## 5. Conclusion

This report documents characteristics of the *F. kawagutii* response to N deficiency, nitrate, and ammonium using physiological, cytometric, and transcriptomic analyses. Our results revealed that nitrogen deficiency caused cell growth depression, cellular cholorophyll *a* depression, lower energy production, lower ability of scavenging ROS, and less resource investment in photosynthate transport. Furthermore, our results also reveal that *F. kawagutii* utilizes ammonium preferentially and faster than nitrate, and nitrate uptake is significantly inhibited by the co-occurrence of ammonium. The molecular mechanism behind the inhibitory effect on nitrate uptake and the preference for ammonium needs to be further investigated. Cytometric and transcriptomic analyses between ammonium and nitrate utilization revealed that growing on ammonium produced similar cell yields as nitrate but with a decreased investment in nutrient transport and assimilation, suggesting reallocation of intracellular resource to other metabolic processes such as antioxidative capability under ammonium condition. We also found that high concentration of ammonium could inhibit the growth of *F. kawagutii*. Taken together, all the results in this study raise a possibility that a coral host can use ammonium to regulate, positively or negatively, the growth of this symbiont. In addition, ten highly and stably expressed genes were screened from 11 treatment condition, which can as candidate reference genes that will be useful for future gene expression studies.

## Author contribution

Senjie Lin and Tangcheng Li conceived the study. Tangcheng Li and Xibei Chen performed the experiments. Tangcheng Li and Senjie Lin wrote the manuscript.

## Declaration of competing interest

All authors declare that there are no conflicts of interest regarding this article.

## Acknowledgement

We wish to thank Xin Lin, Ling Li, Chentao Guo, Liying Yu and Xiaohong Yang of Marine EcoGenomics Laboratory of Xiamen University, China for technical advice and assistance. We thank Zhixiong Huang and Shuh-ji Kao for help with measuring the cellular nitrogen and carbon. We also thank Tao Huang and Lifang Wang for help with measuring the concentration of remaining ammonium and nitrate. This work was supported by the Marine S&T Fund of Shandong Province for Pilot National Laboratory for Marine Science and Technology (Qingdao) (No.2018SDKJ0406-3), and Natural Science Foundation of China grant NSFC 41776116, 31661143029.

## Notes

### Competing Interest Statement

The authors have declared no competing interest.

